# One Health Investigation of SARS-CoV-2 Infection and Seropositivity among Pets in Households with Confirmed Human COVID-19 Cases — Utah and Wisconsin, 2020

**DOI:** 10.1101/2021.04.11.439379

**Authors:** Grace W. Goryoka, Caitlin M. Cossaboom, Radhika Gharpure, Patrick Dawson, Cassandra Tansey, John Rossow, Victoria Mrotz, Jane Rooney, Mia Torchetti, Christina M. Loiacono, Mary L. Killian, Melinda Jenkins-Moore, Ailam Lim, Keith Poulsen, Dan Christensen, Emma Sweet, Dallin Peterson, Anna L. Sangster, Erin L. Young, Kelly F. Oakeson, Dean Taylor, Amanda Price, Tair Kiphibane, Rachel Klos, Darlene Konkle, Sanjib Bhattacharyya, Trivikram Dasu, Victoria T. Chu, Nathaniel M. Lewis, Krista Queen, Jing Zhang, Anna Uehara, Elizabeth A. Dietrich, Suxiang Tong, Hannah L. Kirking, Jeffrey R. Doty, Laura S. Murrell, Jessica R. Spengler, Anne Straily, Ryan Wallace, Casey Barton Behravesh

## Abstract

**Background:** Approximately 67% of U.S. households have pets. Limited data are available on SARS-CoV-2 in pets. We assessed SARS-CoV-2 infection in pet cohabitants as a sub-study of an ongoing COVID-19 household transmission investigation.

**Methods:** Mammalian pets from households with ≥1 person with laboratory-confirmed COVID-19 were eligible for inclusion from April–May 2020. Demographic/exposure information, oropharyngeal, nasal, rectal, and fur swabs, feces, and blood were collected from enrolled pets and tested by rRT-PCR and virus neutralization assays.

**Findings:** We enrolled 37 dogs and 19 cats from 34 of 41 eligible households. All oropharyngeal, nasal, and rectal swabs tested negative by rRT-PCR; one dog’s fur swabs (2%) tested positive by rRT-PCR at the first animal sampling. Among 47 pets with serological results from 30 households, eight (17%) pets (4 dogs, 4 cats) from 6 (20%) households had detectable SARS-CoV-2 neutralizing antibodies. In households with a seropositive pet, the proportion of people with laboratory-confirmed COVID-19 was greater (median 79%; range: 40–100%) compared to households with no seropositive pet (median 37%; range: 13–100%) (p=0.01). Thirty-three pets with serologic results had frequent daily contact (≥1 hour) with the human index patient before the person’s COVID-19 diagnosis. Of these 33 pets, 14 (42%) had decreased contact with the human index patient after diagnosis and none (0%) were seropositive; of the 19 (58%) pets with continued contact, 4 (21%) were seropositive.

**Interpretations:** Seropositive pets likely acquired infection from humans, which may occur more frequently than previously recognized. People with COVID-19 should restrict contact with animals.

**Funding:** Centers for Disease Control and Prevention, U.S. Department of Agriculture

## Introduction

SARS-CoV-2, the cause of the coronavirus disease 2019 (COVID-19) pandemic, likely originated in bats.^1^ Threats from pathogens shared by humans and animals highlight the need for a One Health approach for detection, prevention, and control.^2^ One Health is a collaborative, multisectoral, and transdisciplinary approach with the goal of achieving optimal health outcomes recognizing the interconnection between people, animals, plants, and their shared environment.

In the United States (U.S.), approximately 85 million households (67%) own ≥1 pet, with dogs (63 million households) and cats (43 million households) being most popular.^3^ Human-animal interactions are associated with improved mental, social, and physiologic health^4^ and are critical for people with service and working animals^5^.

Some animals, including pets, have been naturally infected with SARS-CoV-2, almost exclusively after exposure to an infected person.^6–8^ Dogs, cats, ferrets, hamsters, and rabbits are pet species with demonstrated susceptibility to SARS-CoV-2 infection under experimental conditions. Cats, ferrets, and hamsters can transmit the virus to naïve cohabitants of the same species.^9–14^ Additionally, no virus, including SARS-CoV-2, has ever been reported as a contaminant on pet fur. However, animal health and welfare concerns have been reported,^15,16^ including reports of misuse of cleaning products on pets to the Pet Poison Hotline (R. Schmid, personal communication).

We conducted a One Health household transmission investigation to better characterize SARS-CoV-2 infection in mammalian pets living in households with people with COVID-19 to inform guidance and decision-making during this pandemic and for future preparedness efforts.

## Research in context

### Evidence before this study

Both natural and experimental infections with SARS-CoV-2 have been reported in multiple species of companion animals, including dogs, cats, ferrets, hamsters, and rabbits. Cats, ferrets, and hamsters can transmit SARS-CoV-2 to naïve members of the same species. Natural infections of companion animals have occurred almost exclusively after contact with a person with COVID-19.

### Added value of this study

This is one of the earliest studies to assess risk and behavioral factors related to SARS-CoV-2 transmission between people and pets. In households with humans with laboratory-confirmed COVID-19 and pets, 20% had pets with serological evidence of past SARS-CoV-2 infection. SARS-CoV-2 seropositivity in pets was more prevalent among households with higher rate of human COVID-19 infections, and less prevalent among households where owners limited interactions with pets when the owner’s COVID-19 symptoms began. To our knowledge, this is the first study to detect RNA of SARS-CoV-2, or any virus, on animal fur.

### Implications of all the available evidence

Understanding the epidemiologic role that animals may play in the COVID-19 pandemic is crucial to inform guidance and decision making for public health and animal health officials. Our findings add to the growing body of evidence demonstrating SARS-CoV-2 transmission can occur between people and pets—most often from people to pets—and suggest this transmission may occur more frequently than previously recognized. These data highlight the importance of further research, including identification of specific risk factors for human-to-pet transmission, addressing pets in public health guidance during pandemics, and including pets in future pandemic preparedness planning.

## Methods

### Participant Enrollment

The U.S. Centers for Disease Control and Prevention (CDC) collaborated with local and state public health and agriculture departments in Utah and Wisconsin, Wisconsin Veterinary Diagnostic Laboratory (WVDL), and USDA to conduct a One Health investigation that enrolled mammalian pets from an ongoing COVID-19 household transmission investigation that included households with ≥1 person with laboratory-confirmed COVID-19 captured by public health surveillance, previously described.^17^ Human household members with nasopharyngeal or nasal swabs positive by real-time reverse-transcription polymerase chain reaction (rRT-PCR) or who had SARS-CoV-2 antibodies were classified as having laboratory-confirmed COVID-19^17^; additionally, human household patients reporting any symptoms since illness onset of the index case were considered symptomatic. The investigation enrolled human index COVID-19 patients (hereafter addressed as index patients) and household contacts in March 2020 from 62 households to determine secondary household infection rates over a 14-day follow up period since household enrollment. Detailed epidemiologic, clinical, and exposure information was collected for all human household members; most human household members had respiratory specimens collected for SARS-CoV-2 viral testing and blood for serology testing at ≥2 time points. Physical characteristics of each residence, including size, were also described.^17^

Of 62 enrolled households, 41 households with ≥1 mammalian pet living in the household were eligible for inclusion in this One Health investigation (Figure S1). Eligible households were contacted by phone during March–April 2020. Pets were enrolled if owners consented, a questionnaire was completed, and ≥1 sample was collected from each pet. Phone interviews were conducted prior to initial home visits to identify pet species residing in the home and whether the pet(s) developed clinical signs consistent with SARS-CoV-2 infection after the index patient’s COVID-19 diagnosis.

### Household Visits

Initial household visits for pet sampling occurred between April–May 2020 after enrollment in this investigation. Pet sampling was conducted in coordination with repeat visits for the human investigation where possible. During the first household visit for pet sampling, CDC field teams administered a questionnaire (Supplementary Material 1) to capture information on each pet’s demographics, past medical history, household knowledge of public health recommendations, and the following variables for the pet after the index patient’s illness onset: clinical signs; household and community interactions; and household and personal precautionary measures taken. Households were also given an educational information sheet on animals and SARS-CoV-2 (Supplementary Material 2).

During household visits, veterinarians attempted to collect oropharyngeal, nasal, rectal, and fur swabs, feces, and blood from pets. Bilateral deep nasal, oropharyngeal, and rectal swabs were collected, when possible, using sterile polyester tipped swabs (tip diameter, 1.981 mm for nasal, 5.2 mm for oral and rectal). Swabs were placed into 3mL of brain heart infusion broth. Fur swabs were collected in duplicate using 2×2-inch sterile gauze pads rubbed across the back and the abdomen, as well as the dorsal and ventral paws and between the metacarpal and digital pads of each pet. One sample was stored dry and one was stored in RNAlater (Thermo Fisher Scientific, Waltham, Massachusetts). All samples, except for dry fur swabs and fecal samples, were stored on ice packs for immediate shipping and were processed for testing upon arrival at WVDL (Madison, Wisconsin). Dry fur swabs and fecal samples were placed in containers without media and were frozen immediately at −80°C until testing. Serum samples were obtained from venous blood (1–3mL) collected and processed in serum separator tubes; sera were decanted and stored at −80°C until testing.

### rRT-PCR and Serology of Animal Specimens

Preliminary RNA extraction and rRT-PCR testing of animal specimens occurred at WVDL (Supplementary Methods). If rRT-PCR was positive at WVDL for either target, the sample was considered a presumptive positive and sent to the national animal reference laboratory, USDA’s National Veterinary Services Laboratories (NVSL; Ames, Iowa) for confirmatory testing per the USDA Case Definition (https://www.aphis.usda.gov/animal_health/one_health/downloads/SARS-CoV-2-case-definition.pdf). One dry fur swab, the duplicate of the positive fur swab stored in RNAlater, was forwarded to NVSL for confirmatory testing, including rRT-PCR, sequencing, and viral culture attempts (Supplementary Methods). The positive fur swab stored in RNAlater was forwarded to CDC to attempt sequencing (Supplementary Methods). Serum neutralizing antibodies were assessed at NVSL by a SARS-CoV-2 virus neutralization (VN) assay (Supplementary Methods). Neutralizing titers of 1:8–1:16 were considered suspect in the absence of other positive findings; titers > 1:16 were considered seropositive.

### Analysis

Characteristics of enrolled pets, risk factors for seropositivity, number of human cases and household infection rates, and clinical features of human cases within households were analyzed using SAS version 9.4 (SAS Institute, Cary NC). Clopper-Pearson (exact) method was used to calculate 95% confidence intervals for seropositivity rates. Frequent daily contact was defined as having a duration of interaction >1 hour/day between the index patient and the pet (range:1–>12 hours). Features of households with and without seropositive pets were compared using Mann-Whitney-Wilcoxon tests.

## Results

Initial household visits for pet sampling occurred from 0–32 days (median: 14 days) after enrollment in the household transmission investigation. Fifty-six pets (37 dogs, 19 cats) from 34 of 41 eligible (83%) households were enrolled (Figure S1; Table 1); 21 households had only dog(s), seven households had only cat(s), and six households had dogs and cats. Median household size was 4 people (range: 2–8) and 1 pet (range: 1–5) (Table 2). The median proportion of human household members with laboratory-confirmed COVID-19 was 45% (range: 13%–100%); of 72 total people with confirmed infection, 71 (99%) ever experienced symptoms. Additional household characteristics are described in Table 2. Fifty-six pets (100%) had oral and fur swabs, 55 (98%) had nasal swabs, 54 (96%) had rectal swabs, 14 (25%) provided fecal samples, and 47 (84%) provided blood samples. Fourteen pets had repeat oral, nasal, rectal, and fur swabs, 6 had repeat fecal samples, and 11 had repeat blood samples.

**Table 1.**
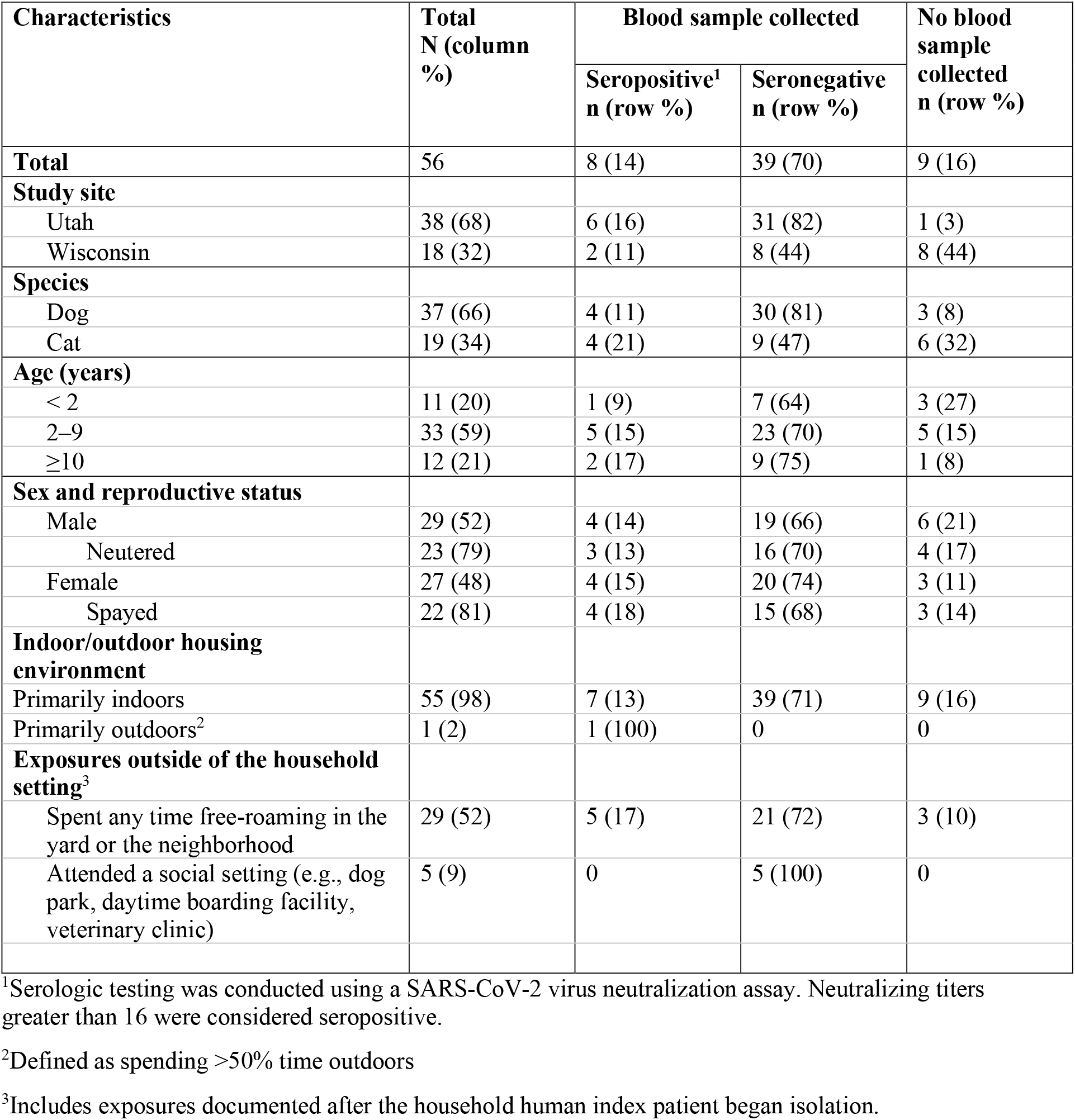
Characteristics of household pets enrolled in the One Health COVID-19 Household Transmission Investigation, April–May 2020.

**Table 2:**
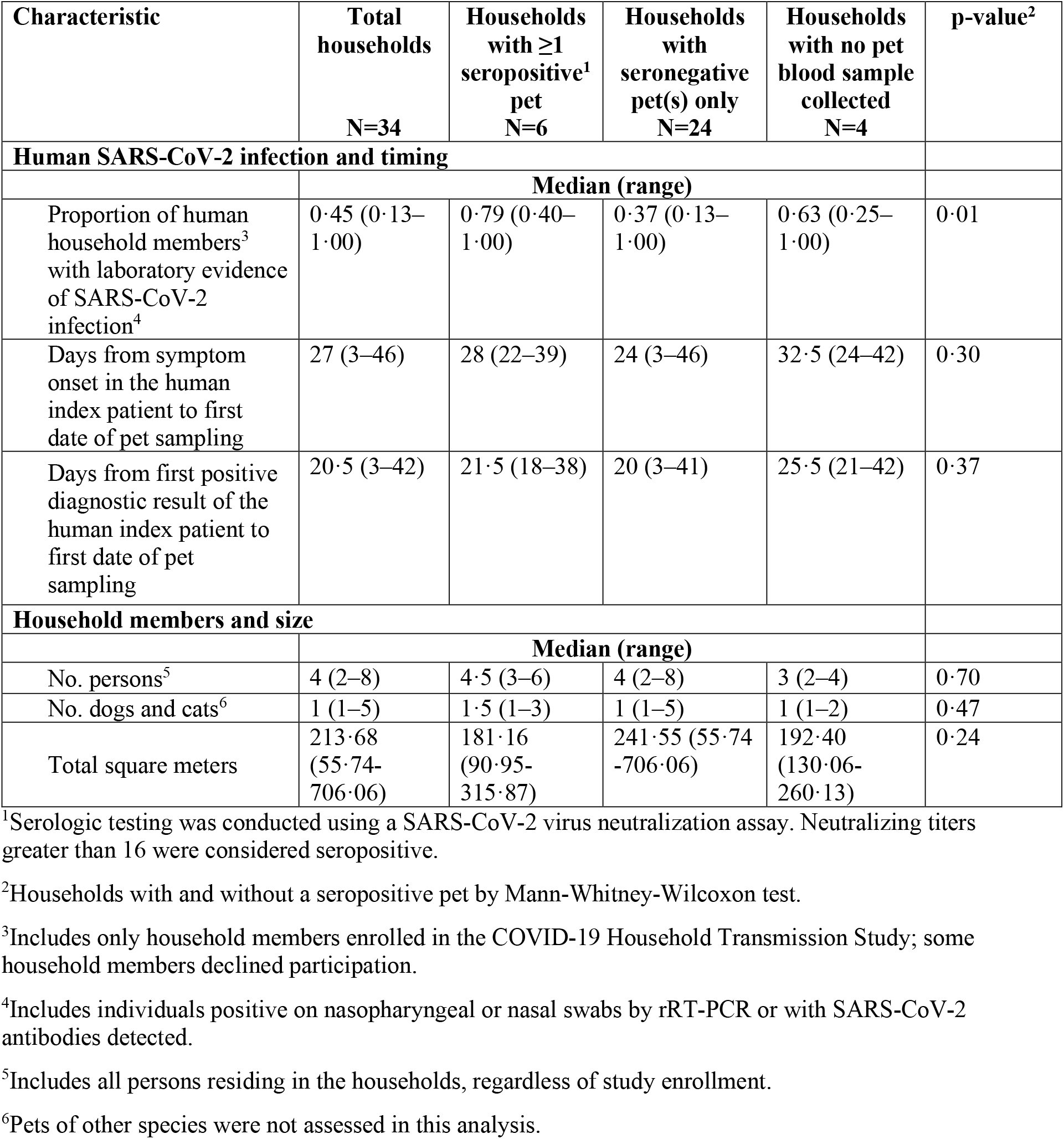
Characteristics of humans with SARS-CoV-2 infection, household members, and timing of human illness in households with pets enrolled in the One Health COVID-19 Household Transmission Investigation, April–May 2020

| Characteristic | Total households<br>N=34 | Households with $\geq 1$ seropositive<sup>1</sup> pet<br>N=6 | Households with seronegative pet(s) only<br>N=24 | Households with no pet blood sample collected<br>N=4 | p-value<sup>2</sup> |
| --- | --- | --- | --- | --- | --- |
| **Human SARS-CoV-2 infection and timing** |  |  |  |  |  |
|  | **Median (range)** |  |  |  |  |
| Proportion of human household members<sup>3</sup> with laboratory evidence of SARS-CoV-2 infection<sup>4</sup> | 0.45 (0.13–1.00) | 0.79 (0.40–1.00) | 0.37 (0.13–1.00) | 0.63 (0.25–1.00) | 0.01 |
| Days from symptom onset in the human index patient to first date of pet sampling | 27 (3–46) | 28 (22–39) | 24 (3–46) | 32.5 (24–42) | 0.30 |
| Days from first positive diagnostic result of the human index patient to first date of pet sampling | 20.5 (3–42) | 21.5 (18–38) | 20 (3–41) | 25.5 (21–42) | 0.37 |
| **Household members and size** |  |  |  |  |  |
|  | **Median (range)** |  |  |  |  |
| No. persons<sup>5</sup> | 4 (2–8) | 4.5 (3–6) | 4 (2–8) | 3 (2–4) | 0.70 |
| No. dogs and cats<sup>6</sup> | 1 (1–5) | 1.5 (1–3) | 1 (1–5) | 1 (1–2) | 0.47 |
| Total square meters | 213.68 (55.74–706.06) | 181.16 (90.95–315.87) | 241.55 (55.74–706.06) | 192.40 (130.06–260.13) | 0.24 |

<sup>1</sup>Serologic testing was conducted using a SARS-CoV-2 virus neutralization assay. Neutralizing titers greater than 16 were considered seropositive.
<sup>2</sup>Households with and without a seropositive pet by Mann-Whitney-Wilcoxon test.
<sup>3</sup>Includes only household members enrolled in the COVID-19 Household Transmission Study; some household members declined participation.
<sup>4</sup>Includes individuals positive on nasopharyngeal or nasal swabs by rRT-PCR or with SARS-CoV-2 antibodies detected.
<sup>5</sup>Includes all persons residing in the households, regardless of study enrollment.
<sup>6</sup>Pets of other species were not assessed in this analysis.

The median time from symptom onset of the index patient to first date of pet sampling was 27 days (range: 3–46 days; Table 2). The median time from first positive diagnostic result of the index patient to first date of pet sampling was 20·5 days (range: 3–42 days) and was similar between households with and without seropositive pets (21·5 vs. 20 days).

All oropharyngeal, nasal, and rectal swabs and fecal specimens tested negative by rRT-PCR, except one rectal swab sample from a cat was presumptive positive that was not confirmed (Supplementary Materials; Table S1). Among 47 pets with serological results from 30 households, eight pets (17%; 4 dogs, 4 cats) from 6 (20%) households, had detectable SARS-CoV-2 neutralizing antibodies. Three pets from these 6 households had seronegative results. The neutralizing titers for all seropositive dog samples were 32 while cat titers ranged from 32 to 128 (Table S1). Demographic pet data by serology result are presented in Table 1. Timelines for human and animal sample collection among households with seropositive pets, as well as symptom onset and duration in people in those households, are depicted in **Figure 1**.

**Figure 1.**
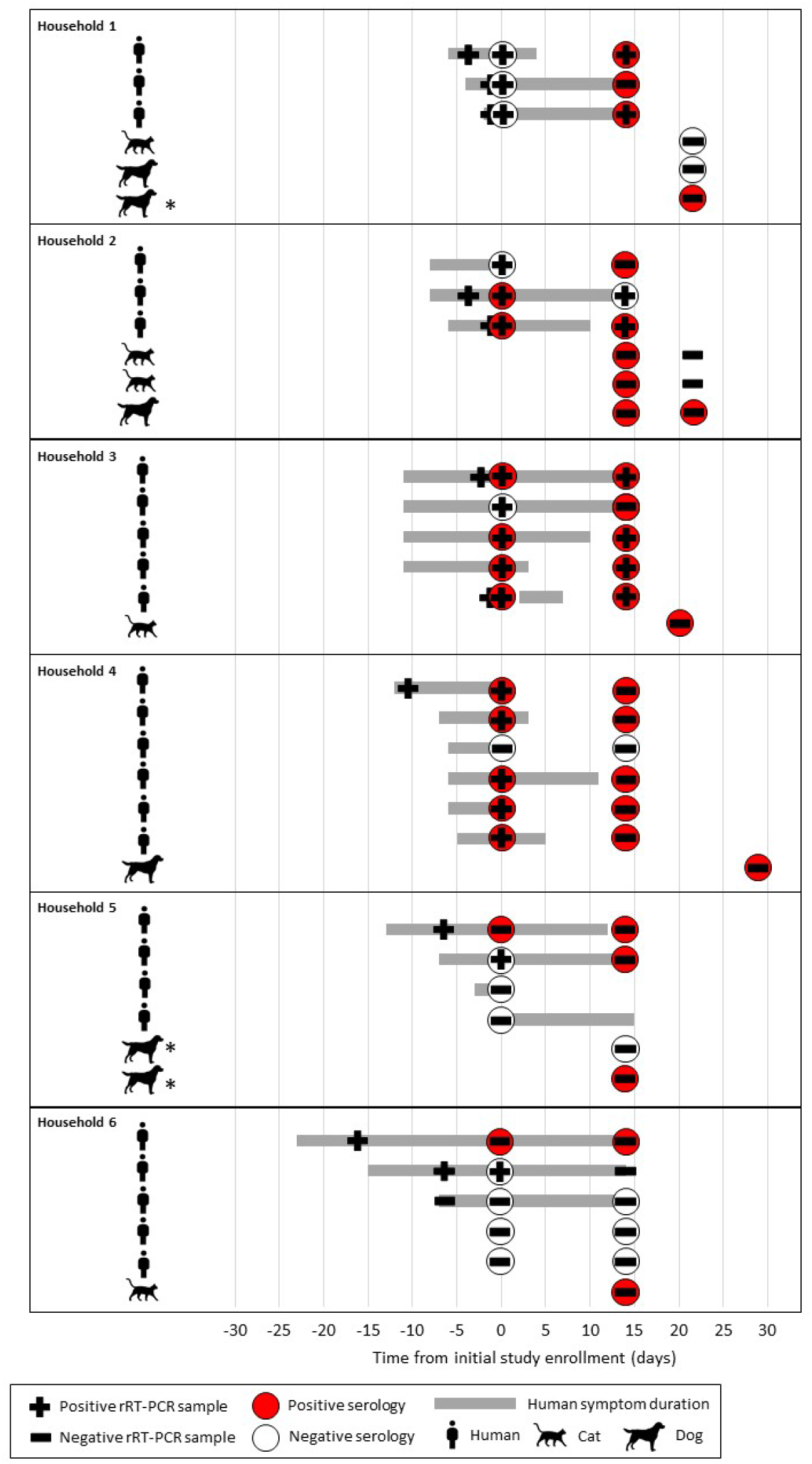
COVID-19 diagnostic testing and symptom duration among humans and animals in households with a seropositive pet, One Health COVID-19 Household Transmission Investigation, April–May 2020. Symptoms durations are shown only for humans. Pets with clinical signs are denoted with an *.

SARS-CoV-2 RNA was detected from duplicate fur swabs from one of 56 pets (2%) at the first pet sampling visit and subsequent fur swabs from this dog were negative (Figure 2). The day the positive fur swab was collected, all six human household members reported symptoms consistent with COVID-19. Five people had nasopharyngeal swabs collected on that day, and four were positive by rRT-PCR. The person who was initially not tested and the one who was initially negative were tested two days later, both were positive. (Figure 2). Seven near-complete or complete-genomes were generated from this household; one each from humans 1–3, three from human 4 collected at three time-points, and one consensus sequence from the dog fur swabs. High sequence similarity suggests one introduction from the community and subsequent internal household transmission (Figure 2; Figure S2). Notably, the dog had no evidence of infection; all samples were negative by rRT-PCR and the dog was also seronegative (Figure 2). Viral culture was attempted on the rRT-PCR positive fur swab, but was negative.

**Figure 2.**
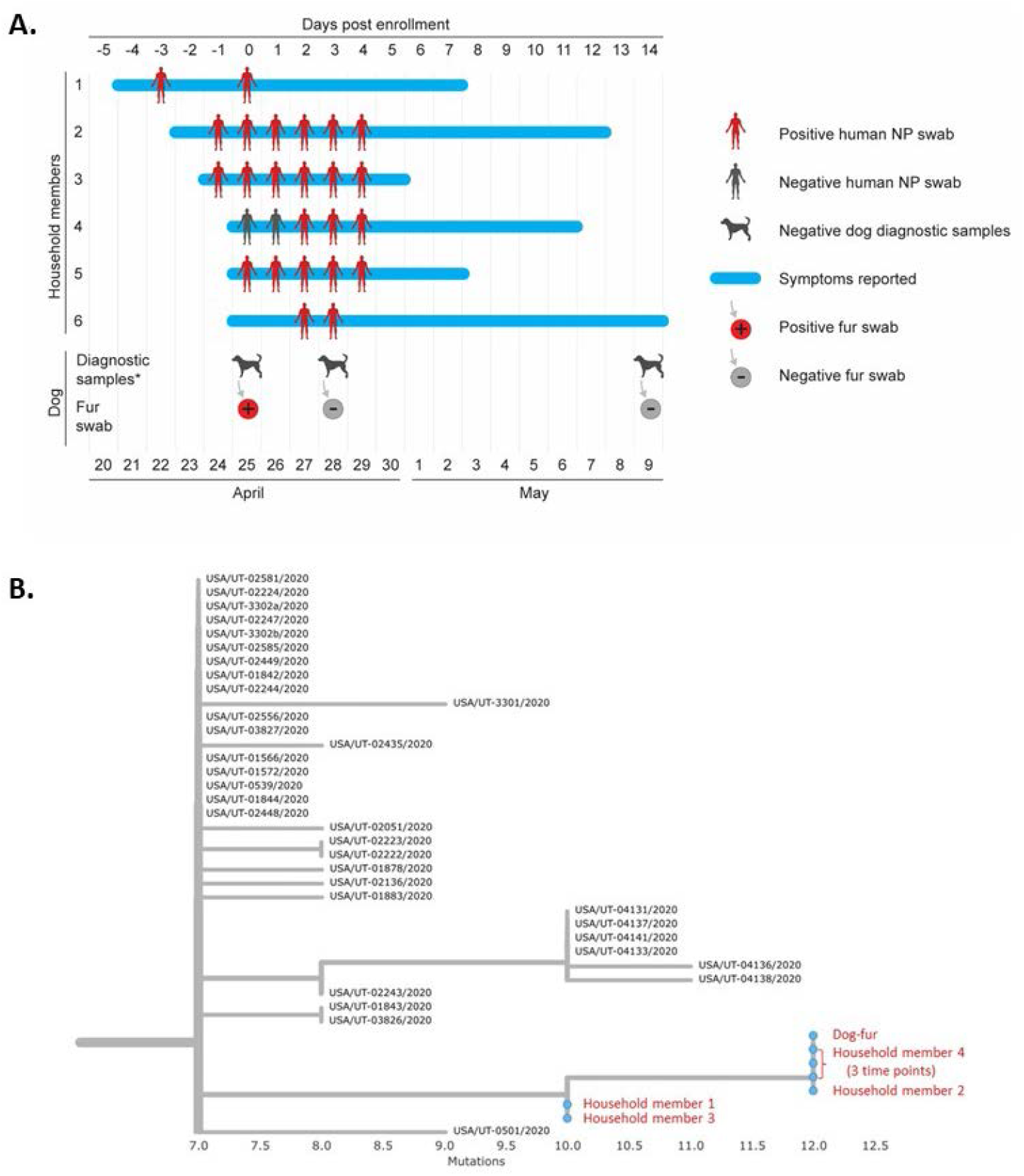
Timeline and phylogenetic analysis of human and dog testing in one household in Utah with SARS-CoV-2 RNA detected on the dog’s fur, One Health COVID-19 Household Transmission Investigation, April–May 2020. Panel A. Timeline of human and dog testing in one household in Utah with six persons with laboratory-confirmed COVID-19 and SARS-CoV-2 RNA detected on the dog’s fur. The timeline indicates dates of reported symptoms and results of nasopharyngeal swab testing by rRT-PCR in human COVID-19 cases and samples collected from the dog in the household. Diagnostic samples* from the dog included oral, nasal, and rectal swabs and stool, which all tested negative by rRT-PCR, and a blood sample which was negative by virus neutralization. Panel B. Enhanced view of branch-tip from comprehensive phylogram (see Figure S2), depicting here the seven study sequences (red) alongside selected Utah complete genome sequences available from Global Initiative on Sharing All Influenza Data. Branch length is by divergence. See Figure S2 for zoomed-out dendrogram depicting additional available sequences from Utah.

Owners reported clinical signs consistent with SARS-CoV-2 infection among 14 (25%) pets during the time from symptom onset of the index patient until time of sampling (Table S2). The most reported clinical signs were respiratory (16%), including sneezing (7%), coughing (7%) and nasal discharge (5%). Among the 8 seropositive pets, clinical signs were reported in only 2 (25%); one dog had nasal discharge and one dog had decreased appetite. Among 39 seronegative pets, clinical signs were reported in 8 (21%) (Table S2).

Forty-six (98%) of 47 pets with serological results were primarily indoor pets; one pet, an 8-year-old seropositive cat, spent ≥50% time outdoors (Table 1). Seropositivity among pets occurred more commonly among households with higher rates of secondary transmission among people; the median proportion of people with laboratory-confirmed COVID-19 in households with a seropositive pet was 79% (range: 40–100%) compared to 37% in households with no seropositive pet (range: 13–100%) (p=0.01) (Table 2). Overall, owners reported pets had fewer daily interactions lasting ≥1 hour and fewer types of interaction with the index patient after their COVID-19 diagnosis; interactions included petting, cuddling, feeding, sleeping in the same location, pets licking the index patient’s face or hands, taking for walks, sharing food, and grooming (Figure 3). Among the 47 pets with serologic results, 33 (70%) pets were reported to have frequent daily contact (≥1 hour) with the index patient before the person’s diagnosis. Of 14 pets with decreased interactions, none (0%) were seropositive. Nineteen pets continued to have frequent contact with the index patient after their diagnosis; of these, 4 (21%) were seropositive. Five (15%) of 34 households, comprising 12 (21%) pets, reported that, after their COVID-19 diagnosis, the index patient began wearing face masks and 2 (6%) also reported glove use around pets. In households using face masks, among pets with serological results, one of eight (13%) pets was seropositive, while in households not using face masks, seven of 39 (18%) pets were seropositive.

Of 34 households, 10 (29%) identified a household member familiar with CDC recommendations for people with suspected or confirmed COVID-19 restricting contact with pets^18^; three (30%) of the 10 households had a seropositive pet. Of the 10 households familiar with CDC recommendations, implementation of precautions was low; the index patient in one (10%) household reduced interactions with pets after the person’s diagnosis, one (10%) household used masks and gloves while interacting with pets, and one (10%) household reported both reduced interaction and mask and glove use.

**Figure 3.**
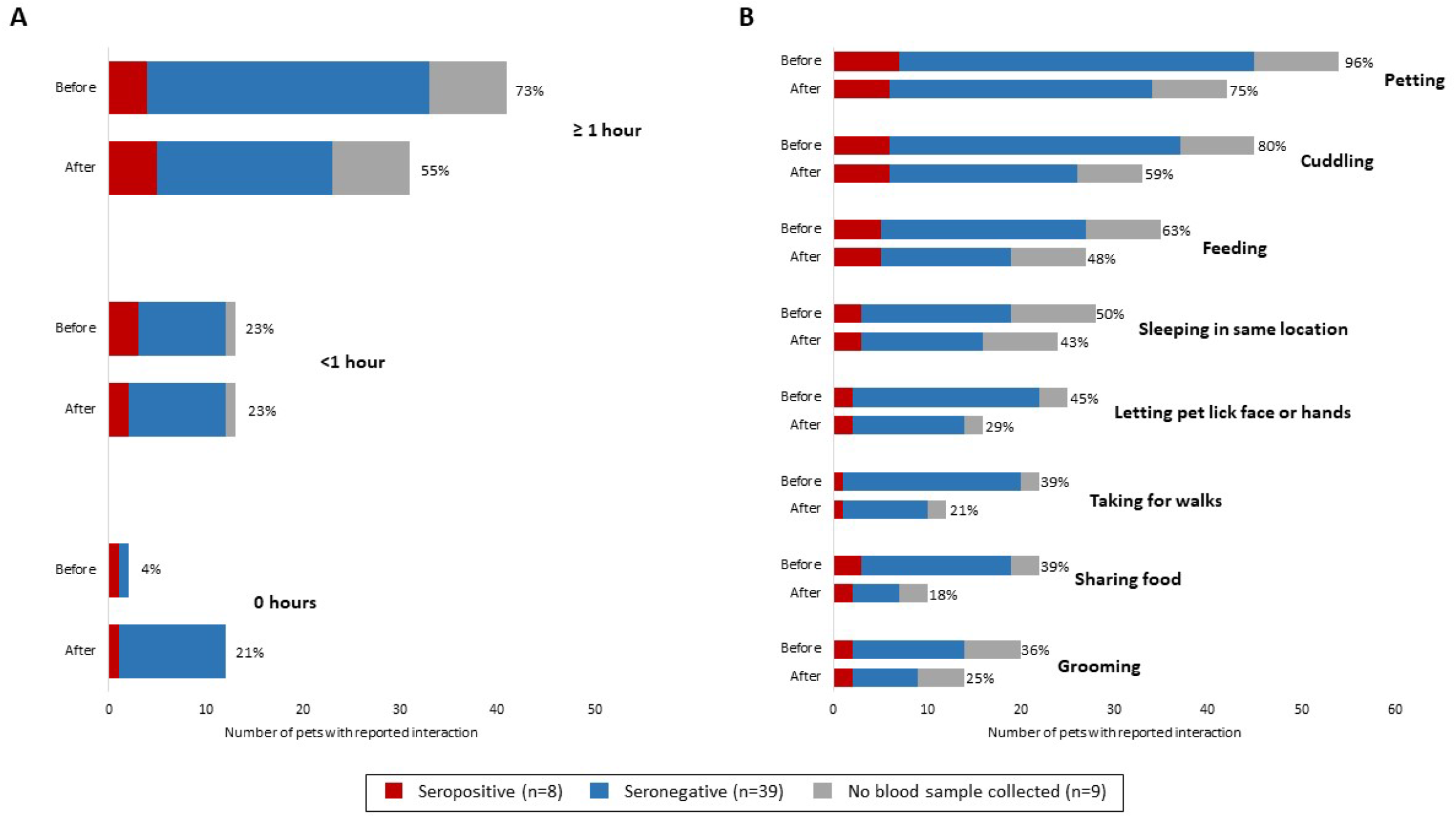
(A) Reported duration of interaction per day and (B) types of interactions between human index patients and pets in each household before and after human index patient diagnosis, by pet serostatus – One Health COVID-19 Household Transmission Investigation, April–May 2020

## Discussion

The epidemiologic role of pets in the COVID-19 pandemic is not fully understood. This One Health investigation systematically evaluated pets residing in households with people with laboratory-confirmed COVID-19. At the time this investigation began, three countries had reported natural SARS-CoV-2 infection in 11 animals, including household pets.^8,19,20^ This investigation identified a higher rate of seropositivity (17%) across enrolled pets with serological results living in households with human COVID-19 cases compared to previously published studies.^7,21,22^ The 12% seropositivity rate in dogs with serological results in our investigation was similar to a previous study^22^; however, the 31% seropositivity in cats is higher than previous reports that range from 0-15% seropositivity.^7,22,23^ While 25% of pets were reported to have clinical signs consistent with SARS-CoV-2 infection, no animals received veterinary treatment specific to these signs. Only two seropositive animals identified were reported to have mild clinical signs consistent with SARS-CoV-2 infection during the period when the infection was most likely. Clinical signs consistent with SARS-CoV-2 infection in animals are generally non-specific and could potentially be attributed to other factors. Cross-species zoonotic transmission events are documented, but are likely under-recognized because of asymptomatic pet infections, small sample sizes, and few published studies with variable results.^21,22^

In our investigation, more seropositive pets were found in households with a greater rate of human household secondary transmission. Further investigations are needed to evaluate SARS-CoV-2 transmission dynamics between people and pets including anthropogenic or mechanical factors such as whether isolation precautions were taken or infectious dose was altered by differences in viral shedding; architectural differences among homes; ventilation system usage patterns affecting air flow; environmental cleaning; or personal protective equipment use. Analysis of household prevention measures, such as facemask use by index patients, was limited by small sample sizes in this study; further investigations are needed to characterize the effectiveness of these measures to prevent SARS-CoV-2 transmission to pets.

Several seropositive animals identified roamed freely in the yard or neighborhood during their likely infectious window, which raises concern for potential transmission of virus from infected pets to people and susceptible animals, which is biologically plausible, but has not yet been documented. One seropositive cat spent ≥50% of its time outdoors. Experimental studies have documented that cats with SARS-CoV-2 can transmit SARS-CoV-2 to other cats,^13,24^ leading to concerns of transmission between cats that roam outdoors; however, this was not assessed in this investigation.

We detected SARS-CoV-2 RNA in fur swabs collected from only one dog but were not able to culture the virus from these samples. Thirty (54%) pets were sampled at a time when at least one household member was symptomatic and 14 (25%) pets at a time when at least one household member tested positive; therefore, some environmental contamination from human viral shedding may have been missed. Our findings suggest that viral RNA on the fur was due to environmental contamination from human household members. Fomite transmission from pet fur seems unlikely although more studies are needed to determine the potential of pet fur to serve as a fomite for SARS-CoV-2 transmission. In households where the index patient decreased duration of interaction with pets after the person’s diagnosis, no pets in this study were seropositive. In two households with seropositive pets, the index patient increased their duration of interaction with pets after their diagnosis (Figure 3). This finding highlights the importance of people with suspected or confirmed COVID-19 restricting contact with pets and other animals to prevent person-to-animal transmission, in accordance with CDC recommendations^25^.

We identified 10 households with awareness of CDC’s recommendations of restricting interactions with pets for people with COVID-19^25^ before enrollment. While this metric was captured only at a single time point, it emphasizes the importance of providing accurate and timely health protection messaging for pets during a pandemic caused by an emerging zoonotic disease.

Our findings provide additional characterization of potential SARS-CoV-2 transmission from people with laboratory-confirmed COVID-19 to pets in households; however, several limitations are noted. While directionality cannot be proven based on these results, the epidemiological information gathered, in conjunction with what is currently known about disease course and shedding of SARS-CoV-2 in companion animals, suggests that human infection preceded animal infection. In experimental infection studies, viral RNA was detected up to the study endpoint--12 days post-infection for cats^9,12,13^, while only on day 6 for dogs^9^. However in cases of natural infection, viral RNA was detected up to 14 and 19 days in dogs^6^ and cats^26,27^, respectively, post-confirmatory testing of the index patient. In this investigation, the median time from symptom onset of the index patient to specimen collection was 27 days (range:3–46 days) and the median time from first positive diagnostic result of the index patient to specimen collection was 20·5 days (range:3-42 days), which would have missed the shedding window for infected pets and could explain the lack of viral RNA detection. The time to pet sampling from the index patient’s symptom onset and from diagnosis were similar among households with and without seropositive pets, and therefore, most pets had a similar length of time to mount neutralizing antibody responses since the beginning of their exposure to the household’s human case(s). Additionally, the sample size of enrolled and tested pets was insufficient to allow for definitive conclusions regarding risk factors for pet infection and to compare interactions between pets and index patients.

Future investigations of household transmission should sample pets across the spectrum of exposure, including time points closer to the start of the index patient’s exposure window and at multiple subsequent time points to learn more about viral shedding, symptomatology, and risk factors. Further One Health efforts are needed to better understand the risk of SARS-CoV-2 transmission between people and pets and to further characterize the course of SARS-CoV-2 infection in pets, both of which will inform guidance and decision-making to best protect public health, animal health, and welfare. This investigation shows that transmission of SARS-CoV-2 from people to pets can occur in household settings. We identified a higher rate of seropositivity than previous studies. Given the relative frequency of human-to-animal transmission in households with people with COVID-19, people with confirmed or suspected COVID-19 should restrict contact with pets and other animals^18^. If a person must care for their pet while they are sick, they should wear a mask and should wash their hands before and after interacting with them^18^.

## Conclusions

A One Health approach for the prevention and control of SARS-CoV-2^2,28^, as well as other emerging and zoonotic diseases, is critical, including response and surveillance efforts to capture and assess transmission dynamics between people, animals, and their shared environment. Previous zoonotic and infectious disease investigations have highlighted the importance of including pets in household transmission investigations. Based on limited information available to date, the risk of pets spreading COVID-19 to people appears low. This One Health investigation provides additional evidence that pets can be infected with SARS-CoV-2, especially after contact with people with COVID-19. Pets contribute to people’s health and well-being, and proper prevention measures to limit microbial transmission between people and pets should be taken to prevent zoonotic infections.

### Data Sharing

Complete or near-complete genome sequences of SARS-CoV-2 obtained in this investigation are available at Global Initiative on Sharing All Influenza Data (GISAID) and GenBank. Additional information or de-identified data may be made available to researchers who submit a methodologically sound proposal to the corresponding author.

## Acknowledgments

The authors wish to thank members of the CDC COVID-19 One Health Working Group, CDC COVID-19 Household Transmission Investigation Field Investigations Teams in Utah and Wisconsin, Drs. Aron Hall, Scott Nabity, Jacqueline Tate, and others on the CDC COVID-19 Epidemiology Task Force. Additionally, we would like to thank all our local, state, and federal human and animal health/laboratory partners. We also thank Hannah Rettler and colleagues from the Utah Department of Health and Dr. Stephen Welch for graphic design assistance with the figures. Lastly, a sincere thank you to the participating households in the Salt Lake City and Milwaukee metropolitan areas for their interest in this investigation and willingness to involve their pets, which made this One Health investigation possible.

## Disclaimers

The findings and conclusions in this report are those of the authors and do not necessarily reflect the official position of the Centers for Disease Control and Prevention or the institutions with which the authors are affiliated.

