## Supplementary Materials for "One Health Investigation of SARS-CoV-2 Infection and Seropositivity among Pets in Households with Confirmed Human COVID-19 Cases — Utah and Wisconsin, 2020"

**Running title**

**Title**

### One Health Investigation of SARS-CoV-2 Infection and Seropositivity among Pets in Households with Confirmed Human COVID-19 Cases — Utah and Wisconsin, 2020

**Authors**

Grace W. Goryoka[1], Caitlin M. Cossaboom[1], Radhika Gharpure[1], Patrick Dawson[1], Cassandra Tansey[1], John Rossow[1], Victoria Mrotz[1], Jane Rooney[2], Mia Torchetti[3], Christina M. Loiacono[3], Mary L. Killian[3], Melinda Jenkins-Moore[3], Ailam Lim[4], Keith Poulsen[4], Dan Christensen[4], Emma Sweet[4], Dallin Peterson[5], Anna L. Sangster[5], Erin L. Young[5], Kelly F. Oakeson[5], Dean Taylor[6], Amanda Price[6], Tair Kiphibane[7], Rachel Klos[8], Darlene Konkle[9], Sanjib Bhattacharyya[10], Trivikram Dasu[10], Victoria T. Chu[1], Nathaniel M. Lewis[1], Krista Queen[1], Jing Zhang[1], Anna Uehara[1], Elizabeth A. Dietrich[1], Suxiang Tong[1], Hannah L. Kirking[1], Jeffrey R. Doty[1], Laura S. Murrell[1], Jessica R. Spengler[1], Anne Straily[1], Ryan Wallace[1], Casey Barton Behravesh[1]

**Author Affiliations**

[1]Centers for Disease Control and Prevention, Atlanta, Georgia (G. Goryoka, C. Cossaboom, R. Gharpure, P. Dawson, J. Rossow, C. Tansey, V. Mrotz, V. Chu, N. Lewis, K. Queen, J. Zhang, A. Uehara, E. Dietrich, S. Tong, H. Kirking, J. Doty, L. Murrell, J. Spengler, A. Straily, R. Wallace, C. Barton Behravesh)

[2]United States Department of Agriculture, Animal and Plant Health Inspection Service, Veterinary Services, Fort Collins, CO (J. Rooney)

[3] United States Department of Agriculture, Animal and Plant Health Inspection Service, Veterinary Services, National Veterinary Services Laboratories, Ames, IA (M. Torchetti, C. Loiacono, M. Killian, M. Jenkins-Moore)

[4]Wisconsin Veterinary Diagnostic Laboratory, Madison, Wisconsin, USA (A. Lim, K. Poulsen, D. Christensen, E. Sweet)

[5]Utah Department of Health, Salt Lake City, Utah, USA (D. Peterson, A. Sangster, E. Young, K. Oakeson)

[6]Utah Department of Agriculture and Food, Salt Lake City, Utah, USA (D. Taylor, A. Price)

[7]Salt Lake County Health Department, Salt Lake City, Utah, USA (T. Kiphibane)

[8]Wisconsin Department of Health Services, Madison, Wisconsin, USA (R. Klos)

[9]Wisconsin Department of Agriculture, Trade and Consumer Protection, Madison, Wisconsin, USA (D. Konkle)

[10]City of Milwaukee Health Department Milwaukee, Wisconsin, USA (S. Bhattacharyya, T. Dasu)

**Corresponding author**

Grace W. Goryoka, 1600 Clifton Rd, MS H16-5; Atlanta, Georgia, 30033; Phone: +1 (404) 718-3466; Fax: +1 (404) 718-1900;

#### Supplementary Materials

**Supplementary Material 1: Animal questionnaire used during the One Health COVID-19 Household Transmission Investigation, April–May 2020**

HHTm
Investigation_Househc

**Supplementary Material 2: Information sheet on animals and SARS-CoV-2 provided to households with pets enrolled in the One Health COVID-19 Household Transmission Investigation, April–May 2020**

One Health COVID19
- Animal information s

### Supplementary Methods

##### RNA Extraction and rRT-PCR of Animal Specimens

Preliminary RNA extraction and real-time reverse-transcription polymerase chain reaction (rRT-PCR) testing of animal specimens occurred at WVDL. To evaluate fecal samples, a swab of feces was obtained and suspended in 1 mL PBS. For all swab specimens, RNA was extracted from 50 µL of sample (in brain heart infusion media or PBS) along with Xeno RNA (10,000 copies, Thermo Fisher Scientific) using the MagMAX-96 Viral RNA Isolation Kit (Thermo Fisher Scientific) on a 96-well KingFisher Flex extraction platform and eluted in a volume of 50 µL according to manufacturer's instructions. 5 µL of extracted RNA was used for a one-step rRT-PCR targeting the 2019-nCoV N gene sequences (N1 and N2). The rRT-PCR assay was based on CDC assay[1] with alternative reagents using the 2019-nCoV RUO Kit (Integrated DNA Technologies, Catalog # 10006713), the AgPath-ID One-Step rRT-PCR Reagents and the VetMAX Xeno Internal Positive Control- LIZ assay (Thermo Fisher Scientific). The rRT-PCR amplification was performed with 1 cycle at 50°C for 15 mins and 90°C for

10 mins, followed by 40 cycles of 95ºC for 15 secs and 55ºC for 1 min on an Applied Biosystems 7500 Fast Real-Time PCR Instrument.

At NVSL RNA was extracted from 50 µL of sample using the MagMAX-96 Viral RNA Isolation Kit (Thermo Fisher Scientific) on a 24-well KingFisher extraction platform and eluted in a volume of 90 µL according to manufacturer's instructions. A modified CDC one-step rRT-PCR N-target assay [N1 and N2 targets][1,2] was used on an Applied Biosystems 7500 Fast Real-Time PCR Instrument according to Emergency Use Authorization instructions for use. Specimens testing presumptive positive at WVDL and confirmed by NVSL were considered SARS-CoV-2–positive. An inconclusive result refers to when only one of two targets (N1 or N2 gene sequences) of the rRT-PCR assay was detected.[1]

**Virus neutralization**

For VN, 25µL of two-fold serially diluted sera (for final dilutions of 1:8 to 1:512) were pre-incubated with 25µL of 100 TCID50/ml of SARS-CoV-2 (2019-nCoV/USA-WA1/2020) in MEM-E containing 200UI/mL penicillin, 200µg/mL streptomycin, 75µg/ml gentamicin sulfate and 6µg/mL Amphotericin B for 60 min at 37°C with 5% $CO_2$. Each serum sample was tested in duplicate in 96-well plates. At one-hour post-infection, 150µl of Vero 76 cells were added to the virus-serum mixtures. The neutralization titers were determined at three days post infection. The titer was recorded as the reciprocal of the highest serum dilution that provided 100% neutralization of the reference virus, as determined by visualization of cytopathic effect. Of approximately 620 sera from cats and dogs, 27% have tested positive by the VN at NVSL. The specificity of the VN assay was assessed in-house by testing sera with antibodies to transmissible gastroenteritis, porcine epidemic diarrhea virus, porcine hemagglutinating encephalomyelitis virus, bovine coronavirus, and Aleutian disease. Additional specificity testing was conducted on a known panel of 47 sera with antibodies to common feline and canine coronaviruses. All of this testing was negative by the VN assay.

**Genome sequencing and analysis**

At CDC, nucleic acid from four rRT-PCR positive specimens (dog fur stored in RNAlater and three specimens from household member 4 from various time points) was extracted and sequenced using the Oxford Nanopore Technologies MinION and Illumina MiSeq following previously published protocols[4] and consensus sequences were generated with Minimap 2.17 and Samtools 1.9. Missing gaps after MinION and MiSeq sequencing were filled by individual rRT-PCR followed by Sanger sequencing. Consensus complete genome sequences were generated using Sequencher 5.4.6.

At the Utah Public Health Laboratory (Salt Lake City, Utah), residual nucleic acid extractions from rRT-PCR positive specimens from household members 1, 2, and 3 were used for input for the ARTIC amplicon sequencing protocol (Josh

Quick 2020. nCoV-2019 sequencing protocol. protocols.io dx.doi.org/10.17504/protocols.io.bdp7i5rn). The amplicons generated were then sequenced on the Illumina MiSeq platform. Consensus sequences were generated using UPHL's custom analysis workflow Cecret (https://github.com/UPHL-BioNGS/Cecret).

RNA sequencing on the dry dog fur swab was performed at NVSL using the Ion AmpliSeq Kit for Chef DL8 and Ion AmpliSeq SARS-CoV-2 Research Panel (Thermo Scientific, Waltham, MA) and sequenced using an Ion 520 chip on the Ion S5 system using the Ion 510™ & Ion 520™ & Ion 530™ Kit. FASTQ files were shared with CDC to include in analysis.

Representative complete genome sequences were downloaded on July 15, 2020 from GISAID, and phylogenetic relations were inferred using maximum likelihood analyses implemented in TreeTime using the Nextstrain pipeline[5].

**Viral culture of fur swab**

Viral culture was performed in Vero cells (ATCC CCL-81) under biosafety level-3 conditions. Cells were cultured in minimum essential medium with Earle's balanced salt solution (MEM-E) with 5% fetal bovine serum (FBS), 25 µg/ml gentamicin sulfate, and 2 µg/ml Amphotericin B (growth media). Cells were seeded in T25 flasks for at least 48h. Specimens were diluted 1:2 in MEM-E containing 200 UI/mL penicillin, 200 µg/mL streptomycin, 75 µg/ml gentamicin sulfate, and 6 µg/mL Amphotericin B. Cells were inoculated with approximately 1·5 ml of the diluted sample and adsorbed for 1 hour at 37°C. Mock-inoculated cells were used as negative controls. After adsorption, cells were washed three times with MEM-E, replacement medium was added, cells were incubated at 37°C and monitored for cytopathic effect (CPE) once daily for up to seven days. Cell cultures with no CPE were frozen, thawed, and subjected to up to two blind passages, with inoculation of fresh cultures with the lysates as described above. Virus isolation was confirmed in the cell cultures by SARS-CoV-2-specific rRT-PCR using the CDC N1 and N2 primer and probe sets.[1]

#### Supplementary Results

**Unique medical histories and clinical signs**

One pet was reported to have an immunocompromising condition; this was a 14-year-old cat with feline immunodeficiency virus (FIV). This pet was negative for SARS-CoV-2 on serologic testing. Two pets were reported to receive immunosuppressive medications (corticosteroids and Janus Kinase inhibitor, respectively) at the time of sampling; both pets were negative on serologic testing.

**Sequence Analysis of Household UT-36 Human Samples and Dog Fur Swab**

Seven near-complete genomes or complete-genomes were generated from household UT-36; three from household member 4, collected at three different time points, three from household members 1-3, and one from the dog's fur swab. High quality sequences were not recovered from household members 5 and 6, and were excluded from analysis. All sequences from household UT-36 formed a unique closely related or identical cluster that nested within many sequences from the United States, including sequences in the major clade containing the S: D614G mutation[6], characteristic of many sequences from United States and Europe (**Figure S2**). Household member 4 was sampled across three time points, yielding identical sequences. Household member 1, the index patient, had an identical sequence to member 3, and was separated from sequences from members 2 and 4 by 2 mutations (**Figure 1**). A nearly complete genome was also generated from a dog fur swab, which was identifical to members 2 and 4 (**Figure** 1). High similarity in sequences suggests a single introduction from the community and internal transmission within the same household, UT-36.

**Presumptive Positive Sample**

One rectal swab from one cat collected 18 days after the first positive test of a human case in the household was presumptive positive (Ct of 36) at WVDL. During confirmatory testing at NVSL, only one of the two targets that define a positive result was detected (SARS-CoV-2 N2 target; Ct 39.6) and sequence could not be attempted. Virus isolation was negative for this rectal swab. SARS-CoV-2 neutralizing antibodies were detected in this cat's blood sample (titer of 128) (**Table S1**).

**Supplementary Figure 1. Enrollment and sampling of household pets in the One Health COVID-19 Household Transmission Investigation, April–May 20**

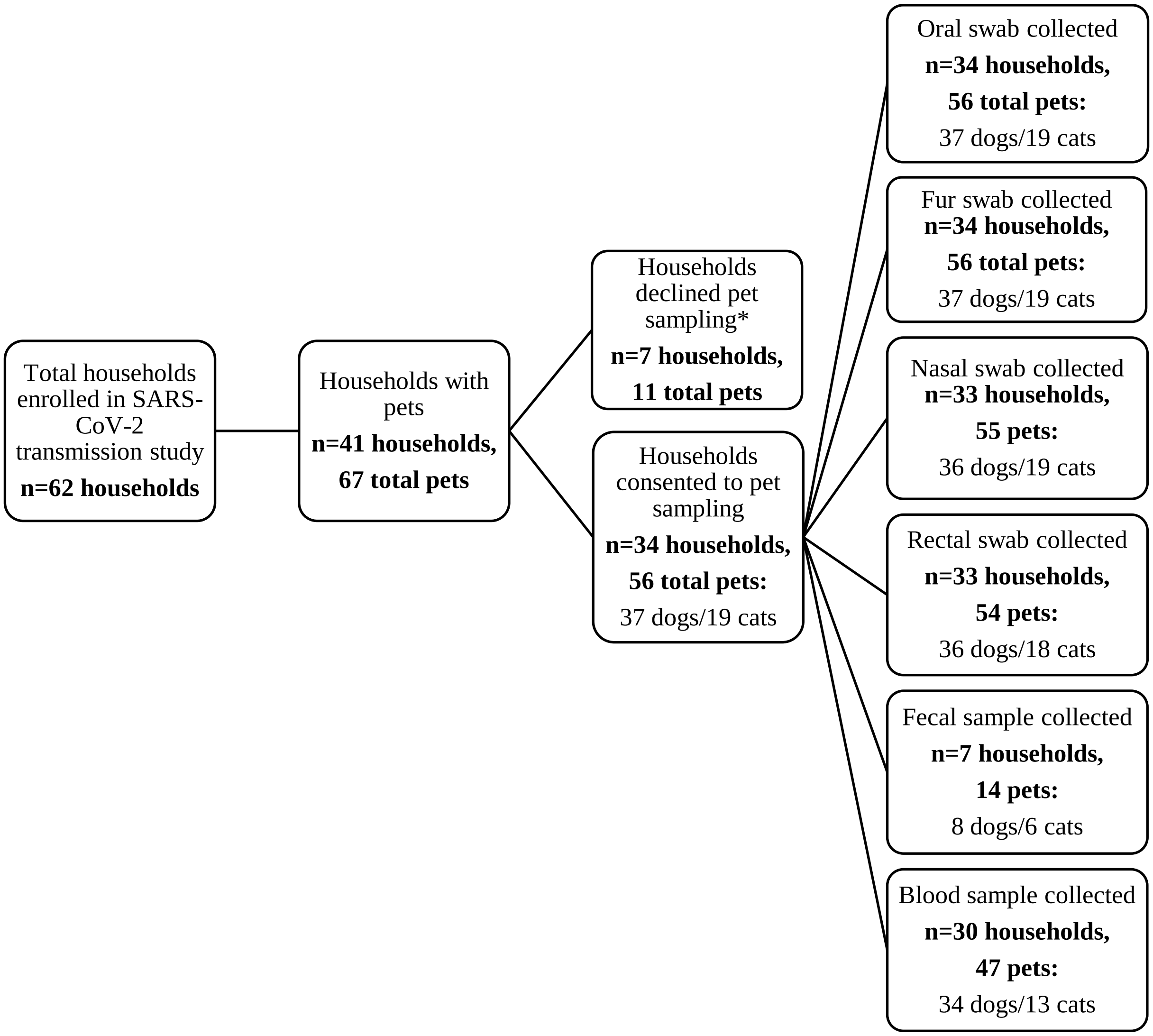


*****Households that declined pet sampling expressed safety concerns for the veterinary field teams due to their pet being fractious.

**Supplementary Figure 2.** Phylogenetic tree with selected Utah complete genome sequences available (as of July 15, 2020) from Global Initiative on Sharing All Influenza Data, color-coded by subclade (19A – 20C); the seven study sequences are shown in red. All study sequences fell within the 20A subclade with the S: D614G mutation. Branch length denotes time interval.

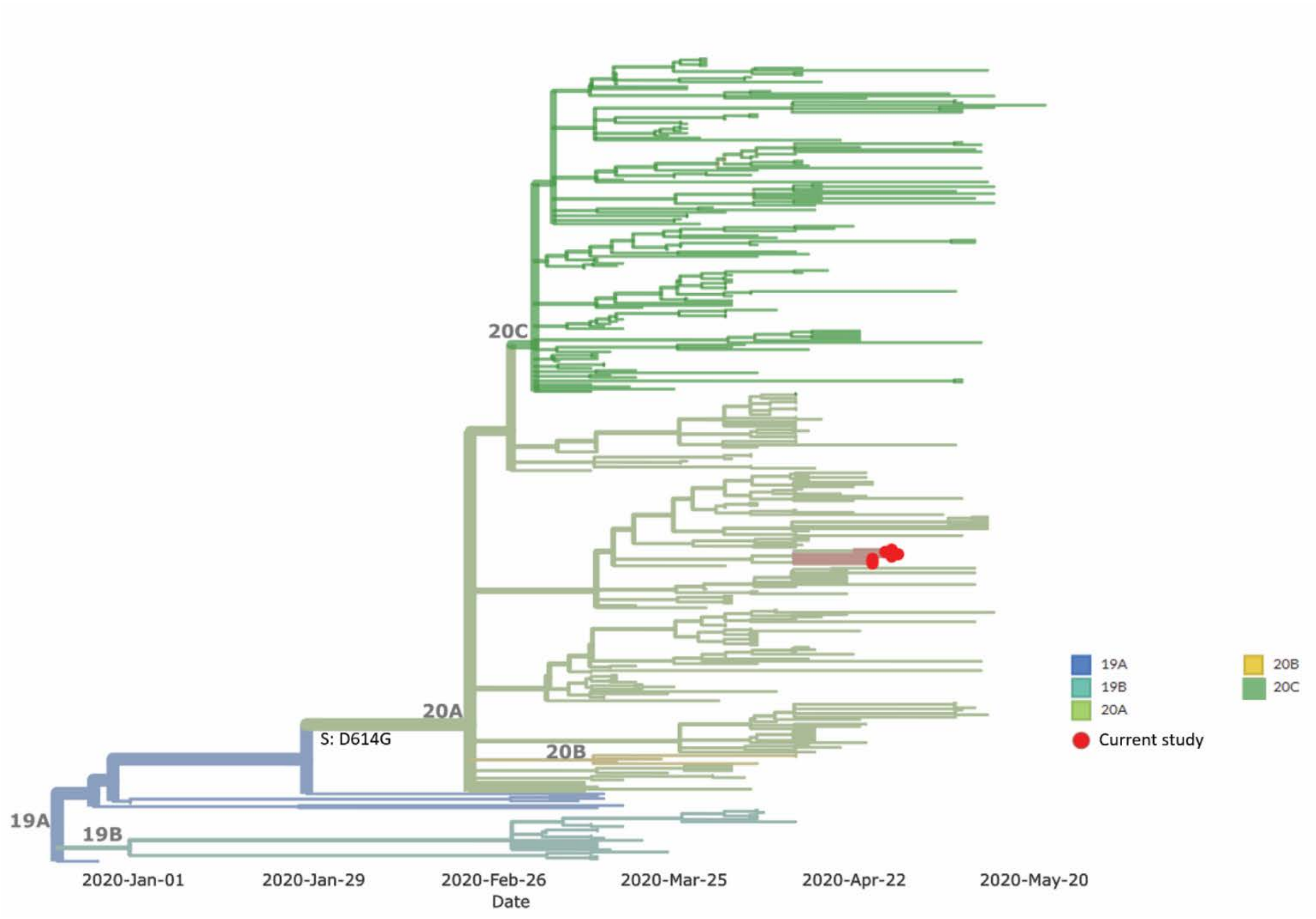

**Supplementary Table 1: Information on seropositive pets and their households from the One Health COVID-19 Household Transmission Investigation, April–May 2020**

| Household | Pet | Species | Sex [*] | Age (years) | # days sample(s) was collected after symptom onset of the household index patient | # days sample(s) was collected after household enrollment | Serostatus (days post-enrollment) | Pre-Existing Condition | Clinical signs[†] | Notes |
|---|---|---|---|---|---|---|---|---|---|---|
| 1 | Dog | Australian Shepherd/ Labrador Retriever mix | MC | 2 | 27 | 21 | Seropositive (1:32) | - | Decreased appetite | All three human household members had at least one positive rRT-PCR result on nasopharyngeal swabs and were seropositive. |
| | Dog | German Shepherd | FS | 12 | 27 | 21 | Seronegative | - | - | |
| | Cat | Domestic shorthair | FS | 14 | 27 | 21 | Seronegative | - | - | |
| 2 | Cat | Domestic longhair | MC | 6 | 22, 29[+], 33 | 14, 21[+], 25 | Seropositive (1:64) on day 14 and 25 | - | - | All three human household members had at least one positive rRT-PCR result on nasopharyngeal swabs and were seropositive. One rectal swab from the 6-year-old female cat was presumptive positive but was not confirmed at NVSL and was negative by virus isolation. |
| | Cat | Domestic medium-hair | FS | 6 | 22, 29[+], 33 | 14, 21[+], 25 | Seropositive (1:128) on day 14 and 25 | Prediabetic; history of asthma | - | |
| | Dog | Miniature Schnauzer | FS | 12 | 22 and 29 | 14 and 21 | Seropositive (1:32) on day 14 and 21 | - | - | |
| 3 | Cat | Domestic shorthair | MC | 8 | 31 | 20 | Seropositive (1:64) | - | - | All five human household members had at least one positive rRT-PCR result on nasopharyngeal swabs and were seropositive. |
| 4 | Dog | Golden Retriever/Pood le mix | MC | 10 | 39 | 29 | Seropositive (1:32) | 15-year history of seizures | - | Four of five human household members had at least one positive rRT-PCR result on nasopharyngeal or nasal swabs and were seropositive; one was seronegative without a positive swab result. |
| 5 | Dog | Pitbull | FS | 3 | 27 | 14 | Seropositive (1:32) | - | Nasal discharge | Two of four human household members had at least one positive rRT-PCR result on nasopharyngeal or nasal swabs and were seropositive; two were negative on initial swabs and serology, and convalescent specimens were not obtained. |
| | Dog | Pitbull | MC | 1 | 27 | 14 | Seronegative | - | Nasal discharge | |
| 6 | Cat | Manx | FS | 8 | 37 | 14 | Seropositive (1:64) | | | This cat spent at least 50% time outdoors. Only one of five human household members was seropositive; an additional member had positive rRT-PCR results on nasopharyngeal swabs but did not have a convalescent blood specimen. |

[*]MC = Male, castrated; FS = Female, spayed
[†]Clinical signs were reported since onset of illness in the first human case in the household. The duration of clinical signs was unknown.
[+]Blood samples were unable to be obtained for these pets on day 21

**Supplementary Table 2. Owner-reported clinical signs among household pets enrolled in the COVID-19 Household Transmission Investigation since onset of illness in first household human case, April–May 2020.**

| Clinical signs | Total<br>N=56<br>n (%) | Seropositive<br>N=8<br>n (%) | Seronegative<br>N=39<br>n (%) | No blood sample collected<br>N=9<br>n (%) |
|---|---|---|---|---|
| None | 42 (75) | 6 (75) | 31 (79) | 5 (55) |
| Any signs reported | 14 (25) | 2 (25) | 8 (21) | 4 (44) |
| Respiratory | 9 (16) | 1 (13) | 6 (15) | 3 (33) |
| Sneezing | 4 (7) | 0 (0) | 2 (5) | 2 (22) |
| Coughing | 4 (7) | 0 (0) | 2 (5) | 2 (22) |
| Nasal discharge | 3 (5) | 1 (13) | 2 (5) | 0 (0) |
| Difficulty breathing / shortness of breath | 1 (2) | 0 (0) | 1 (3) | 0 (0) |
| Gastrointestinal | 3 (5) | 0 (0) | 3 (8) | 0 (0) |
| Vomiting | 1 (2) | 0 (0) | 1 (3) | 0 (0) |
| Diarrhea | 2 (4) | 0 (0) | 2 (5) | 0 (0) |
| Other | 2 (4) | 1 (13) | 0 (0) | 1 (11) |
| Inappetence | 1 (2) | 1 (13) | 0 (0) | 0 (0) |
| Lethargy | 1 (2) | 0 (0) | 0 (0) | 1 (11) |
